# Metabolic control of germ layer proportions through regulation of Nodal and Wnt signalling

**DOI:** 10.1101/2023.12.04.569862

**Authors:** Kristina S. Stapornwongkul, Elisa Hahn, Laura Salamo Palau, Krisztina Arato, Nicola Gritti, Kerim Anlas, Patryk Poliński, Mireia Osuna Lopez, Miki Eibisuya, Vikas Trivedi

## Abstract

During embryonic development, cells exit pluripotency to give rise to the three germ layers. Metabolic pathways influence cell fate decisions by modulating the epigenetic, transcriptional, and signalling states of cells. However, the interplay between metabolism and the major signalling pathways that drive the emergence of ectoderm, mesoderm, and endoderm remains poorly understood. Here, we demonstrate an instructive role of glycolytic activity in activating signalling pathways involved in mesoderm and endoderm induction. Using an in vitro model system for mouse gastrulation, we observed that inhibiting glycolysis prevents the upregulation of primitive streak markers, resulting in a significant increase in ectodermal cell fates at the expense of mesodermal and endodermal lineages. We demonstrate that this relationship is dose-dependent, enabling metabolic control of germ layer proportions through exogenous glucose levels. Mechanistically, we found that glycolysis inhibition leads to the downregulation of Wnt, Nodal, and Fgf signalling. Notably, this metabolic phenotype was rescued by Nodal or Wnt signalling agonists in the absence of glycolytic activity, suggesting that glycolytic activity acts upstream of both signalling pathways. Our work underscores the dependence of specific signalling pathways on metabolic conditions and provides mechanistic insight into the nutritional regulation of cell fate decision making.

## Introduction

Research in the field of stem cell biology has been critical in uncovering the mechanisms that underlie the intricate interplay between metabolism and cell fate determination [1, 2]. It is now widely acknowledged that metabolic pathways not only fulfil the bioenergetic needs of cells but also act as regulators of differentiation. The underlying mechanisms range from metabolite-driven post-translational modifications and metabolite-protein interactions to moonlighting metabolic enzymes, and can affect the epigenetic as well as the signalling state of cells [3–6]. This perspective has gained support from *in vivo* studies that underscore the mechanistic role of metabolism during embryonic development [7–14]. Regulation of cellular metabolic state has been exploited to enhance the efficiency of differentiation and reprogramming protocols [15]. Therefore, a comprehensive understanding of the complex interactions between metabolism, signalling and differentiation will open new avenues for engineering reproducible tissue patterns *in vitro*. In a broader context such an approach will guide our efforts to study the effects of genetic metabolic disorders, malnutrition and maternal diabetes on embryonic development.

One of the earliest cell fate decisions is the exit from pluripotency resulting in the emergence of the three germ layers: the ectoderm, mesoderm and endoderm. In the embryo, prospective mesodermal and endodermal cells migrate through a transient structure known as the primitive streak (PS) while remaining epiblast cells will adapt an ectodermal cell fate [16]. Much attention has focused on how a cell’s preference for either aerobic glycolysis or oxidative phosphorylation changes during differentiation [15]. Pluripotent stem cells (PSCs) are thought to rely on high glycolytic activity to maintain their characteristic histone acetylation patterns [17]. While some studies have suggested that metabolic switching is a prerequisite for epigenetic remodelling and differentiation [17,18], others have found that a shift towards oxidative metabolism is germ-layer specific and only occurs in mesodermal and endodermal cells [19]. In contrast to these findings from directed differentiation of cells using extrinsic signals, a study in the developing tailbud (neuromesodermal progenitors, NMPs) found that inhibition of glycolysis increased the proportion of neuroectoderm at the expense of the presomitic mesoderm via regulation of Wnt signalling [14]. Unlike the bipotent NMPs, cells of the early embryo can still give rise to all future cell types. Therefore, the relationship between metabolism and signalling during the earliest stages of differentiation of pluripotent embryonic cells into the three germ layers remains unresolved. Recent studies, both *in vitro* and *in vivo*, have reported spatiotemporal restriction of different glucose transporters that accompany germ layer patterning and how metabolism affects signalling particularly during mesoderm specification [20, 21]. However, the simultaneous control of the three germ layers and their cell type-specific interplay between glycolysis and signalling remains obscure.

In this study, we further elucidate the interplay between glycolysis and the signalling pathways that coordinate germ layer differentiation. Using 3D mouse gastruloids, a stem-cell based model system that allows the co-differentiation of the three germ layers [22], we found that glycolysis plays a crucial role in both endoderm and mesoderm induction by activating Nodal, Wnt and Fgf signalling. Importantly, exogenous glucose (Glc) concentration has a dose-dependent effect on PS marker expression and the development of endodermal and mesodermal cell type derivatives. Thus, we show for the first time that glycolytic activity is not merely permissive but rather acts as an instructive signal which can be used to control germ layer proportions. Moreover, we were able to decouple the metabolic phenotype of glycolysis inhibition from its effects on gastruloid development by rescuing mesoderm and endoderm induction with agonists of the Nodal and Wnt signalling pathways. This demonstrates that glycolytic activity is not a bioenergetic prerequisite for endoderm and mesoderm induction, but instead functions as an important activator of Nodal and Wnt signalling. These findings demonstrate how metabolic activity acts as a regulator of morphogen signalling and cell fate determination and opens new possibilities for metabolic control of cell type proportions in *in vitro* systems.

## Results

### Glycolysis is needed for T/Bra expression and symmetry breaking in gastruloids

To address the role of metabolism during germ layer specification, we used gastruloids, aggregates of mouse embryonic stem cells (mESCs) which specify cell types of all three germ layers while establishing an anterior-posterior (AP) axis [22]. Between 24 and 48 hours post aggregation (hpa), gastruloids upregulate the PS and early mesoderm marker Brachyury (T/Bra) throughout the tissue (Figure 1A) [23,24]. In a symmetry-breaking event, the posterior pole is then specified by the polarisation of T/Bra expression. At 72hpa, markers of all three germ layers can be detected and further culture results in gastruloid elongation along the AP axis [25]. In contrast to previous culture protocols, we did not add the glycogen synthase kinase-3 (GSK3) inhibitor and Wnt signalling activator CHIR99021 (CHIR) to provide a clean delineation between signalling and metabolic activity. Even in the absence of CHIR treatment, we find that 95% of gastruloids generated from E14 T/Bra::GFP mESCs break spontaneously symmetry and elongate when they were previously maintained in serum/LIF conditions (Figure 1A).

**Fig. 1.**
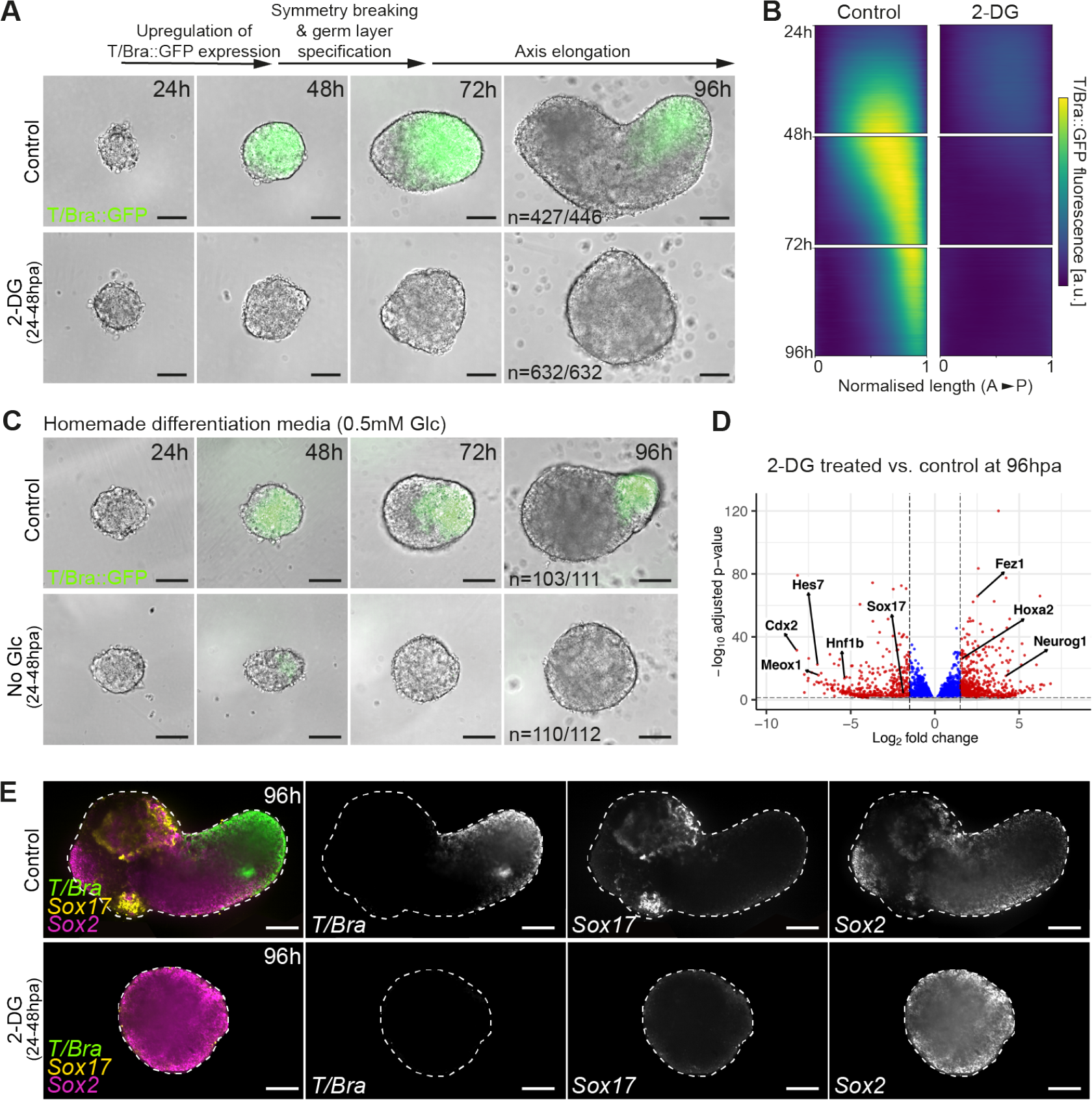
Glycolysis is required for T/Bra expression, mesoderm and endoderm induction. (A) In control gastruloids, T/Bra::GFP reporter expression is first present throughout gastruloid and then gets polarised before gastruloid elongation. Glycolysis inhibition between 24-48hpa (6mM 2-DG) results in loss of T/Bra::GFP expression and impaired gastruloid development. N numbers from *N*_*exp*_ = 12. Scale bars: 100*µ*m. (B) Kymographs of control and 2-DG treated gastruloids. Polarisation of T/Bra::GFP fluorescence in controls while 2-DG treated gastruloids fail to upregulate T/Bra::GFP expression. Normalised length from anterior (A) to posterior (P). Kymographs were generated by averaging T/Bra::GFP fluorescence of *n* = 30 gastruloids for each condition. (C) Gastruloids in homemade differentiation media containing 0.5mM Glc express T/Bra::GFP and break symmetry. Removing Glc between 24-48hpa results in loss of T/Bra::GFP expression. N numbers from *N*_*exp*_ = 3. Scale bars: 100*µ*m. (D) Volcano plot describing differentially expressed genes between 2-DG treated and control gastruloids at 96hpa based on RNA-seq. Red dots describe significantly differentially expressed genes that pass thresholds for log2FC = 1.5 (vertical lines) and adjusted *p* value =0.05 (horizontal line). *n* = 6 for both conditions. (E) HCR stainings of control and 2-DG treated gastruloids at 96hpa. Single confocal slices are shown. *N*_*exp*_ = 3, *n* = 107 (control), *n* = 104 (2-DG). Scale bars: 100*µ*m.

To investigate the role of Glc metabolism during germ layer specification, we used 2-Deoxy-D-glucose (2-DG) to inhibit glycolysis in gastruloids (Figure S1A). Using a T/Bra::GFP reporter line, we found that 2-DG treatment between 24 and 48hpa resulted in the loss of T/Bra::GFP expression and subsequent failure of symmetry breaking and axis elongation (Figure 1A). Time lapse imaging showed upregulation of T/Bra::GFP expression in control gastruloids between 24 and 48hpa, while 2-DG treated gastruloids did not start expressing the PS marker even after 2-DG withdrawal (Figure 1B and Movie S1,2). It has been reported that the effects of 2-DG are not limited to the inhibition of glycolysis but could also affect other cellular functions due to breakdown products of 2-DG [26]. To test whether the phenotype is specific to Glc metabolism, we prepared N2B27 differentiation medium with 0.5mM Glc, reflecting physiological Glc levels in the uterine fluid [27]. While this is around 40 times less than the Glc concentration used in commercially available N2B27 medium, it not only supported T/Bra::GFP expression and symmetry breaking sufficiently, but also allowed to remove Glc between 24 and 48hpa of gastruloid development by repeated washes with Glc-free medium. Similar to the 2-DG treatment, lack of Glc in this time window resulted in reduced T/Bra::GFP upregulation and failure to polarise, suggesting that the observed 2-DG phenotype was indeed mediated by reduced glycolytic activity (Figure 1C). These experiments suggest that active glycolysis is needed during early gastruloid development to ensure T/Bra expression and subsequent symmetry breaking. To test if loss of T/Bra::GFP is specific to glycolysis inhibition, we next treated gastruloids with sodium azide (NaN3), a Complex IV inhibitor that blocks oxidative phosphorylation (OxPhos) (Figure S1A). While the treatment resulted in an effective block of OxPhos, as measured by loss of oxygen consumption as well as reduced gastruloid growth, we did not observe loss of T/Bra::GFP expression and symmetry breaking (Figure S1B,C). These results suggest that metabolic inhibition and reduced cell proliferation does not affect T/Bra::GFP expression per se and indicates that glycolysis might play a specific role in gastruloid development.

### 2-DG suppresses mesoderm and endoderm formation

Gastruloids display post-occipital cell types of all three germ layers which are spatially organised along an AP axis [28]. We next asked how the early inhibition of glycolysis affects cell fate decision making during gastruloid development. To assess the presence of different cell types, we performed single-gastruloid RNA-seq on control and 2-DG treated gastruloids at 96hpa. We found that markers of mesodermal derivatives, such as *Hes7* and *Meox1* were strongly downregulated (Figure 1D, Figure S2A) [29, 30]. Similarly, endodermal markers, such as *Sox17* and *Hnf1b* were reduced in 2-DG treated gastruloids (Figure 1D, Figure S2B) [31]. In contrast, transcripts associated with neural cell fates, such as *Neurog1* and *Fez1*, were upregulated (Figure 1D, Figure S2C) [32, 33]. Furthermore, we found an upregulation of the most anterior Hox gene and hindbrain marker *Hoxa2* while posterior Hox genes and *Cdx2*, which marks the posterior embryo, were strongly reduced (Figure 1D, Figure S2E) [34, 35]. This suggests an anteriorisation of 2-DG treated gastruloids.The shift in germ layer proportions was further confirmed by in situ hybridisation chain reaction (HCR) showing that in contrast to control gastruloids, cells expressing *T/Bra* (PS/early mesoderm) and *Sox17* (definitive endoderm) transcripts were strongly reduced in 2-DG treated gastruloids (Figure 1E). Instead, *Sox2* (ectoderm/pluripotent) was expressed throughout the glycolysis inhibited gastruloids. Further stainings showed that the *Sox2*^+^ domain was subdivided in clusters of cells expressing the neuroectodermal marker *Sox1* or the pluripotency marker *Nanog* (Figure S2F) [36, 37]. Both of these markers were overrepresented compared to control gastruloids, confirming the transcriptomics results (Figure S2C,D). These results show that the inhibition of glycolysis between 24 and 48hpa has long-term effects on gastruloid development and results in a shift of germ layer proportions away from posterior mesodermal and endodermal derivatives towards more anterior neuroectodermal cell fates.

### Glucose has a dose-dependent effect on the proportion of T/Bra::GFP expressing cells

The observed effects of 2-DG treatment on gastruloid development raise the question whether glycolytic activity has a permissive or instructive role in T/Bra expression and germ layer induction. A permissive role would suggest the existence of a certain threshold of cellular glycolytic activity needed to allow T/Bra upregulation in gastruloids. If instead the glycolytic activity acted as an instructive signal, one would expect a dose-dependent effect on T/Bra gene expression. To address this question, we aimed to modulate the glycolytic activity in gastruloids to assess the effect on T/Bra::GFP expression and germ layer proportions. Taking advantage of our *in vitro* system, we varied exogenous Glc levels in the differentiation medium and found that this changes glycolytic activity effectively in 2D cultured cells (Figure 2A). In the absence of Glc, cells relied entirely on OxPhos for ATP production (Figure S3A). Raising Glc levels resulted in increased glycolytic activity which plateaued around 12.5mM Glc. At this concentration, glycolysis and OxPhos contributed approximately evenly to the total ATP production rate (Figure S3A). To test whether Glc and glycolytic activity have a dose-dependent effect on T/Bra::GFP expression, we generated gastruloids in medium containing different amounts of Glc. At 48hpa, when T/Bra::GFP expression is present throughout the gastruloid, a positive correlation between Glc concentration and GFP signal was observed (Figure 2B). However, it was also apparent that gastruloids grown in lower Glc concentrations were strongly reduced in size suggesting that Glc was growth limiting.

**Fig. 2.**
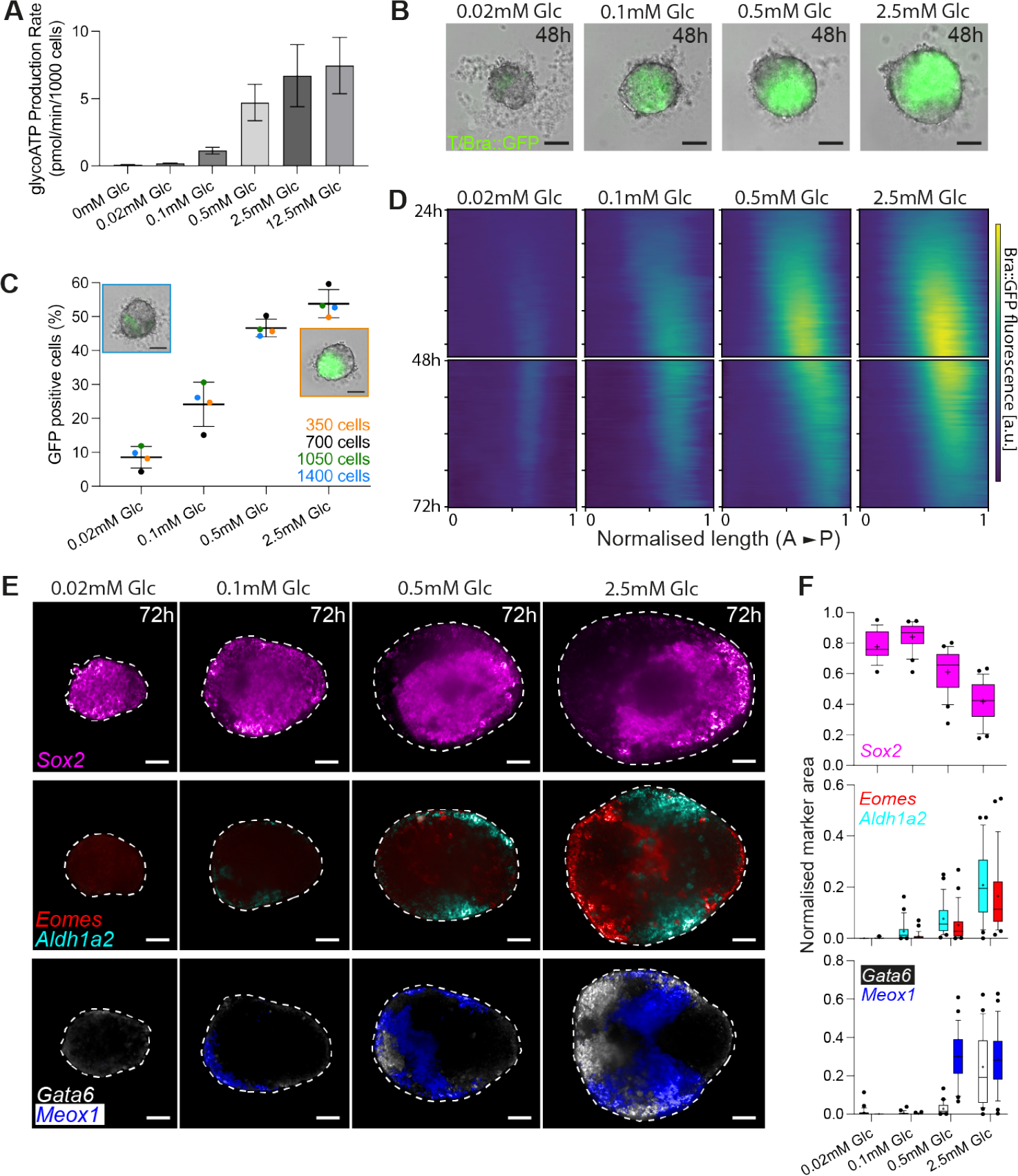
Glucose has a dose-dependent effect on T/Bra::GFP expressing cells and germ layer proportions. (A) Changes in mean glycoATP production rate depending on exogenous Glc levels. Error bars indicate SD. *N*_*exp*_ = 3 for all conditions. (B) Raising exogenous Glc concentration results in increasing levels of T/Bra expression in gastruloids. *N*_*exp*_ = 7, *n* ≥ *315 for each condition. Scale bars: 100*µ*m. (C) Mean percentage of GFP^+^ cells in T/Bra::GFP gastruloids at 48hpa which were generated by aggregating 350, 700, 1050 and 1400 cells and cultured in different Glc concentrations. Error bars indicate SD. Inlets show images of gastruloids generated by aggregation of 1400 (blue frame) and 350 (orange frame) initial cells and grown in 0.02mM and 2.5mM Glc respectively. At 48hpa, gastruloids are similar sized but express different levels of T/Bra::GFP. Scale bars: 100*µ*m. (D) Kymographs of T/Bra::GFP intensity in gastruloids developing in different Glc concentrations from 24 to 72hpa. Kymographs were generated by averaging *n* = 10 gastruloids for each condition. (E) Germ layer marker expression in gastruloids at 72hpa which developed at different Glc concentrations. Single confocal slices are shown. *n* = 3, *N*_*exp*_ ≥ 31 for each condition. Scale bars: 50*µ*m.(F) Marker area quantifications on average Z-projections of the batch of gastruloids shown in (E). Marker area was normalised to gastruloid area. Sox2: *n* = 18 (0.02mM Glc), *n* = 22 (0.1mM Glc), *n* = 23 (0.5mM Glc), *n* = 22 (2.5mM Glc). Eomes/Aldh1a2: *n* = 18 (0.02mM Glc), *n* = 26 (0.1mM Glc), *n* = 25 (0.5mM Glc), *n* = 25 (2.5mM Glc). Gata6/Meox1: *n* = 10 (0.02mM Glc), *n* = 18 (0.1mM Glc), *n* = 22 (0.5mM Glc), *n* = 20 (2.5mM Glc).

To exclude the possibility that size and not glycolytic activity is affecting T/Bra::GFP expression, we generated gastruloids from different initial cell numbers resulting in similar sized gastruloids at different Glc concentrations (Figure 2C, see insets). By assessing the percentage of GFP^+^ cells using flow cytometry at 0.02mM Glc, 0.1mM Glc, 0.5mM Glc and 2.5mM Glc over four different initial cell numbers (350, 700, 1050 and 1400 cells), we found that the percentage of GFP^+^ cells was indeed dependent on Glc concentration and not gastruloid size (Figure 2C, Figure S3B). We further wondered if Glc concentration not only determines the percentage of T/Bra::GFP expressing cells but also cellular T/Bra::GFP expression levels. Based on flow cytometry histograms, we detected no noticeable shift in typical GFP peak intensity levels suggesting that cellular T/Bra::GFP expression levels are similar in GFP^+^ cells across different Glc concentrations (Figure S3C). Exogenous Glc concentration thus determines the likelihood of a cell to switch into a *T/Bra* expressing state in a dose-dependent manner. As a result, gastruloids developing in higher Glc concentration display a higher percentage of T/Bra::GFP at 48hpf, resulting in a higher overall T/Bra::GFP intensity level. These results indicate that glycolytic activity plays an instructive role in *T/Bra* induction during gastruloid development, especially between 0mM Glc and 0.5mM Glc where dose-dependency is most prominent (Figure S3B). Interestingly, Glc concentration in the uterine fluid has been measured to be around 0.6mM [27]. Thus, it is conceivable that the *in vivo* embryo develops in a regime where Glc can act as an instructive regulator of PS induction.

### Glucose has a dose-dependent effect on germ layer proportions

Next, we were wondering whether the dose-dependent effect of Glc on the relative number of T/Bra::GFP expressing cells further translates into a change of the proportion of cell types deriving from the different germ layers. We first determined the dynamics of T/Bra::GFP reporter activity at different Glc concentrations by performing time lapse imaging. Kymographs of T/Bra::GFP reporter intensity showed that gastruloids developing at 0.02mM Glc failed to polarise (Figure 2D). Instead a weak stripe of T/Bra::GFP expression formed, a phenotype previously described in gastruloids treated with Nodal or Fgf signalling inhibitors [25]. Despite differences in reporter intensities, the dynamics of symmetry breaking at 0.1mM, 0.5mM and 2.5mM Glc were comparable (Figure 2D). Next, we performed HCR stainings for *Sox2* and markers of PS derived cell fates. We found that the proportion of *Sox2* expressing cells decreased with rising Glc concentration (Figure 2E,F). In contrast, mesodermal and endodermal markers, such as *Eomes* (endoderm/anterior mesoderm), *Gata6* (cardiac mesoderm/endoderm), *Aldh1a2* (trunk mesoderm) and *Meox1* (paraxial mesoderm) were absent in gastruloids cultured at 0.02mM Glc but showed an increase in proportion with rising Glc concentration (Figure 2E,F). Our results show for the first time that exogenous Glc levels can bias the emergence of specific cell types deriving from the three germ layers in a dose-dependent manner. Thus, increasing Glc levels lead to a greater proportion of mesodermal and endodermal cell derivatives, accompanied by lower proportions of ectodermal cell fates. This demonstrates how glucose concentration can be used as tool to modulate the proportions of cell lineages in gastruloids.

### Nodal, Wnt and Fgf signalling pathway activity is dependent on glycolysis

We next wanted to understand the mechanism by which glycolytic activity regulates changes in cell fate decision making. PS markers are induced and maintained by major developmental signalling pathways, such as Nodal, Wnt and Fgf signalling [38–41]. To test whether the effect of glycolysis inhibition on mesoderm and endoderm development is mediated by reduced signalling, we performed single-gastruloid RNA-seq on control and 2-DG treated gastruloids immediately at the end of the treatment period at 48hpa of gastruloid development. We found that transcription of Nodal, Wnt3a and Fgf8 ligands, as well as many of their target genes, such as *T/Bra, Eomes* and *Dusp6* were downregulated upon 24 hours of glycolysis inhibition (Figure 3A). Among the most upregulated transcripts were *Meis2, Irx3* and *Chrdl1*, all genes associated with neural development (Figure S4A) [42–44]. These results suggest that the effect of reduced glycolytic activity on germ layer proportions can indeed be explained by the reduced activity of Nodal, Wnt and Fgf signalling.

**Fig. 3.**
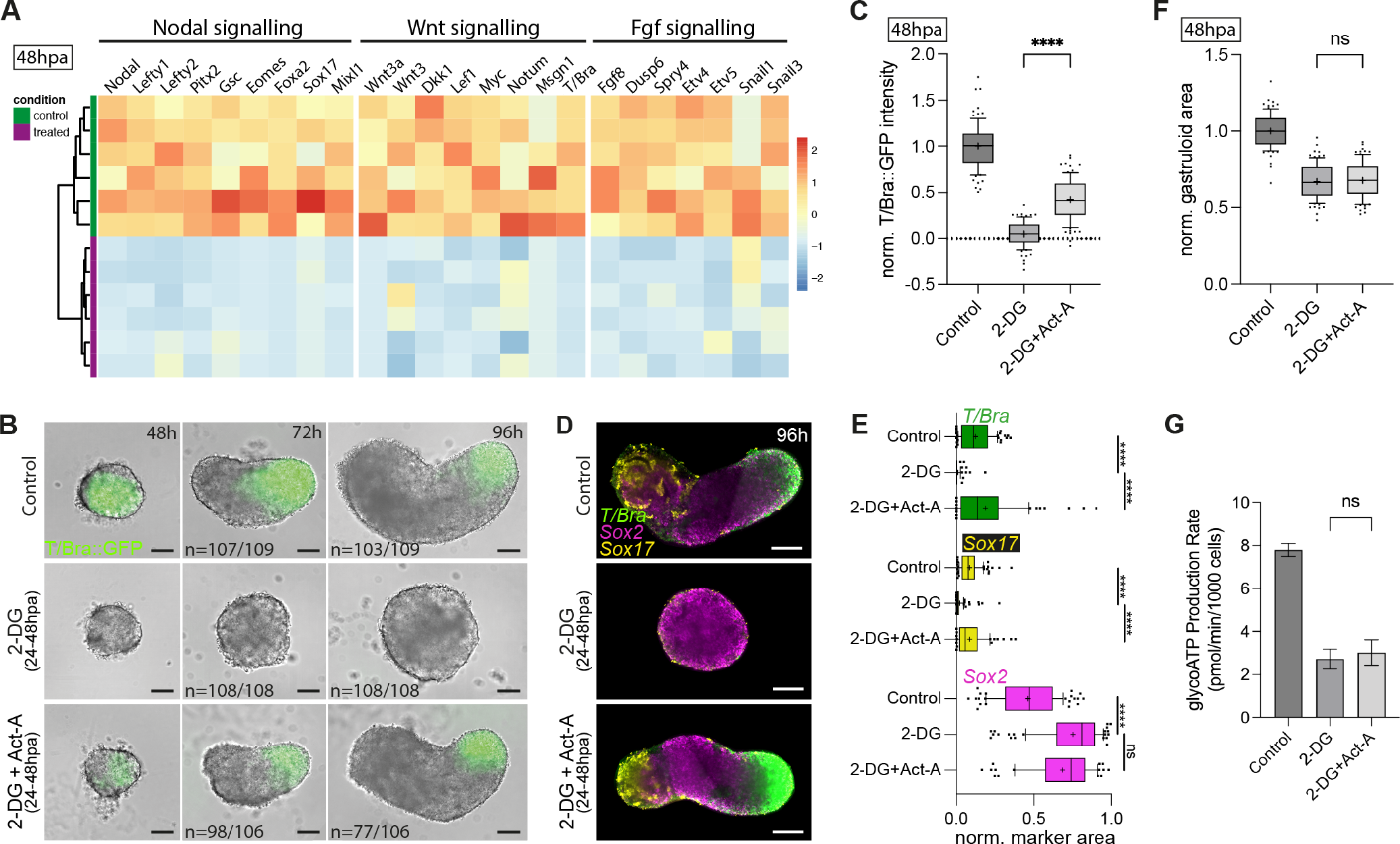
Nodal signalling activation can rescue the 2-DG phenotype in the absence of glycolytic activity. (A) Heatmap depicting relative transcript levels of selected genes in 2-DG treated vs. control gastruloids at 48hpa, based on RNA-seq. List was manually curated based on typical genes for selected signalling pathways. Heatmap columns are normalised per gene. Lower and higher expression in blue and red, respectively. Each row represents a single gastruloid with applied hierarchical clustering. (B) Gastruloids developing under control conditions, glycolysis inhibition or Nodal signalling rescue condition. N numbers for symmetry breaking at 48hpa and axis elongation at 96hpa from *N*_*exp*_ = 4. Scale bars: 100*µ*m. (C) T/Bra::GFP intensity of gastruloids at 48hpa for different treatment conditions. Background was subtracted and intensities were normalised to mean control intensity. *N*_*exp*_ = 3, *n* = 87 (control), *n* = 85 (2-DG), *n* = 87 (2-DG+Act-A). \*\*\*\**p <* 0.0001 for t-test. (D) Images of HCR stainings for control, 2-DG and Nodal signalling rescue. T/Bra: PS/early mesoderm (green), Sox2: ectoderm/pluripotent (magenta), Sox17: definitive endoderm (yellow). Single confocal slices are shown. Scale bars: 100*µ*m. (E) Area quantifications of germ layer markers using average Z-projections of HCR stainings as shown in (D). Area was normalised to total gastruloid area. Line in box plot indicates median, + indicates mean, whiskers indicate 10 to 90 percentiles. *N*_*exp*_ = 3, *n* = 107 (control), *n* = 104 (2-DG), *n* = 70 (2-DG+Act-A). \*\*\*\**p <* 0.0001 for t-test. (F) Gastruloid area at 48hpa for different treatment conditions. Area was normalised to mean control area. *N*_*exp*_ = 3, *n* = 87 (control), *n* = 85 (2-DG), *n* = 87 (2-DG+Act-A). t-test was not significant (ns). (G) Mean glycoATP production rate of cells cultured for 48h in differentiation medium. Inhibitor and growth factor treatment was started 24h prior to measurements. *N*_*exp*_ = 3, error bars indicate SD. t-test was not significant (ns).

### Development of glycolysis inhibited gastruloids can be rescued by Nodal or Wnt signalling pathway activation

The observation that Nodal, Wnt and Fgf signalling pathways are downregulated upon glycolysis inhibition raises the question whether glycolytic activity acts as an activator of signalling pathways or whether signalling is just not functional under the bioenergetic constraints related to the lack of glycolytic activity. If glycolysis is solely required to activate these pathways or to promote cells required to induce these signalling pathways, it should be possible to rescue the DG phenotype by activating signalling pathways during the treatment period. Indeed, simultaneous activation of the Nodal signalling pathway with Activin A (Act-A) during 2-DG treatment between 24 and 48hpa, was able to partially rescue T/Bra::GFP expression at 48hpa (Figure 3B,C). Even though T/Bra::GFP expression was not reaching control levels at 48hpa, symmetry breaking was robust in these rescued gastruloids and elongation was observed in 70% of the cases. We further performed HCR stainings at 96hpa to validate whether cells of all three germ layers can be found in the rescued gastruloids. Quantifications of the relative area of T/Bra and Sox17 expression confirmed that mesoderm and endoderm development was rescued (Figure 3D,E). Further stainings also confirmed the presence of paraxial mesoderm (Meox1) and cardiac precursors (Gata6) in these gastruloids (Figure S4B). These results confirm that Nodal signalling activation during glycolysis inhibition was sufficient to rescue the development of mesodermal and endodermal cell types. Activation of Wnt signalling using CHIR was also able to rescue gastruloid development, albeit in a less reproducible manner (Figure S4C,D). Even though elongation occurred in 34% of gastruloids, endodermal and cardiac cell derivatives were mostly absent, most likely because Wnt signalling strongly promotes differentiation towards posterior mesoderm (Figure S4B,E). We also added Fgf8 simultaneously with 2-DG but did not observe a rescue of T/Bra::GFP induction and polarisation (Figure S4F). Our rescue experiments suggest that Nodal and Wnt signalling activation are downstream of glycolytic activity and can rescue the 2-DG mediated phenotype on gastruloid development.

### Glycolytic activity is not rescued by Nodal and Wnt signalling

The signalling, epigenetic and metabolic state of cells are highly integrated through several feedback mechanisms that ensure coordination of cellular behaviour [1, 4, 6, 45]. In cancer cells, Wnt signalling is a crucial regulator of metabolic reprogramming resulting in increased Glc uptake and its preferential fermentation to lactate even in the presence of oxygen, known as the Warburg effect [46]. Similarly, Wnt, Fgf and Nodal signalling have been suggested to promote glycolysis in the context of normal embryonic development and homeostasis as well as in cancerous tissue [13, 47–50].

To rule out the possibility that Nodal or Wnt signalling activation rescue the 2-DG mediated glycolysis inhibition, we first analysed gastruloid size that is expected to be affected by 2-DG treatment [51]. Indeed, we detected a significant reduction in gastruloid size upon the 24h 2-DG treatment period. In contrast to T/Bra::GFP intensity levels, neither Nodal nor Wnt signalling activation rescued the gastruloid size phenotype suggesting that anabolic metabolism driving proliferation and tissue growth was not restored in these gastruloids (Figure 3F; Figure S4G). To look at a more direct readout of metabolic state, we next measured the ATP production rate from glycolysis (glycoATP production rate). Control cells cultured for two days in differentiation medium displayed a significantly higher glycoATP production rate than cells that had been treated with 2-DG for 24h prior measurement (Figure 3G). We found that the addition of Nodal signalling agonist Act-A did not rescue the glycoATP production rate of 2-DG treated cells. This was also the case when Act-A and CHIR were added simultaneously to cells (Figure S4H). These results were further supported by the finding that T/Bra::GFP expression and symmetry breaking of gastruloids in Glc-free medium (24-48hpa) was also rescued by the addition of Act-A and CHIR (Figure S3I). Here, the absence of Glc abolishes any possibility of reactivating glycolytic activity. Together, these experiments suggest that even though signalling activation can rescue gastruloid development, it does not rescue the metabolic phenotype on the level of glycolytic activity in 2-DG treated cells. This implies that glycolytic activity is not needed for the activation of T/Bra::GFP expression as long as signalling pathways are active. Therefore, glycolysis activity is a crucial inducer of mesodermal and endodermal cell fates but does solely act through the activation of signalling rather than posing bioenergetic constraints on the specification of certain cell types.

## Discussion

### An instructive role of glycolytic activity for mesoderm and endoderm induction

Spatial metabolic patterns observed during amphibian gastrulation in the 1930s sparked early hypotheses about the metabolic regulation of embryonic development [52–54].

In recent years, both *in vitro* and *in vivo* studies have contributed to our understanding of the mechanistic link between metabolic pathways and cell fate specification [4, 55]. Here, we found an important role of glycolytic activity in the induction of endoderm and mesoderm using both a glycolysis inhibitor and Glc-free medium. Most importantly, we demonstrate that this relationship is dose-dependent. As glycolytic activity increases, so does the probability of cells committing to mesodermal and endodermal cell fates. This suggests that glycolysis acts as an instructive cue, rather than merely a permissive factor.

Differences in our findings compared to a prior study [19], which indicated that cells differentiating into endoderm and mesoderm required a shift from high glycolysis to high Ox-Phos, might be attributed to the use of directed differentiation protocols involving growth factor-containing germ layer-specific media. In our gastruloid system, cells of all three germ layers co-emerge without the need for exogenous signalling agonists. This allowed us to demonstrate the importance of gly-colytic activity in the induction of mesoderm and endoderm via Wnt signalling and Nodal signalling activation. Since agonists of these signalling pathways are part of the directed differentiation protocol, it is conceivable that the importance of glycolytic activity for the activation of these pathways could not be detected previously. Our findings align with another recent study that showed adverse effects of 2-DG treatment on mesoderm specification in gastruloids, highlighting the reproducibility of this phenotype [21]. Contrary to our results, the authors did, however, not observe a phenotype in Glc-free medium and concluded that glycolytic activity is dispensable for mesoderm development in gastruloids. A possible explanation might be that the experiments were conducted in the presence of CHIR, a condition that is analogous to our rescue experiments. Since even low amounts of Glc are sufficient for Bra induction, residual Glc after washing out high-Glc differentiation media might be another potential explanation for the observed differences. In accordance with our findings, an important role of glycolysis has been suggested for mesoderm development at later stages of embryonic development when bipotent NMPs commit to either a neural tube or presomitic mesoderm fate [13]. In NMPs it is however still disputed whether glycolytic activity has a promoting or inhibitory function on Wnt and Fgf signalling [12, 14]. Concurrently with our research, another preprint has identified metabolic activity as a key driver of phenotypic variation through integrated molecular-phenotypic profiling of trunk-like structures, an *in vitro* model system for neural tube and somite formation [56]. Specifically, they demonstrate that an early imbalance in OxPhos and glycolysis leads to aberrant morphology and biases cells towards the neural lineage, consistent with our findings during germ layer formation. While the relationship between glycolysis and mesoderm development has been explored in several studies, little is known about the metabolic control of endoderm specification. Previous studies have also found that glycolysis can promote endoderm differentiation [57, 58]. One suggested mechanism involves the Lin41 protein kinase as a non-canonical phosphorylation target of glycolytic enzyme Pfkp, resulting in the suppression of Sox2 and leading to increased endodermal differentiation of ESCs [57]. Here, we show that glycolytic activity is needed for the activation of endoderm-promoting Nodal signalling and therefore introduce an additional layer of metabolic control to our understanding of endoderm specification.

### Bioenergetic versus signalling function of glycolysis

Metabolism plays a vital role in generating the necessary building blocks and energy required for growth and proliferation. Differentiating cells might display unique bioenergetic demands which must be met to ensure the proper development of certain cell types [59, 60]. For instance, the energetic requirements of prospective mesodermal and endodermal cells migrating through the PS might differ significantly from epithelial cells fated to become neuroectoderm. However, our finding that the metabolic phenotype of glycolysis inhibition can be rescued through the activation of Nodal and Wnt signalling pathways in the absence of glycolytic activity strongly indicates that the phenotype is not related to energetic constraints. Since glycolysis is not reactivated in these rescued gastruloids, it seems to primarily act as an activator of developmental signalling pathways. This implies that glycolytic activity is not an absolute necessity for mesoderm and endoderm differentiation as long as these signalling pathways are activated to coordinate gastruloid development by inducing T/Bra upregulation between 24-48hpa. Since morphogenetic movements in the gastruloid model system are not comparable with *in vivo* gastrulation, it is however conceivable that a bioenergetic function of glycolysis plays a role in the migration of mesodermal and endodermal cells [7, 20].

An open question is how changes in glycolytic activity can regulate Nodal, Wnt and Fgf signalling during germ layer induction. Metabolite-driven posttranslational modifications, moonlighting glycolytic enzymes and metabolite-protein interactions are possible links between metabolism and signalling [4, 6, 61]. For instance, high glycolytic activity promotes acetyl-CoA levels which can be rate-limiting for protein acetylation [62]. Interestingly, the activity of Nodal and Wnt signalling transducers is known to be modulated by acetylation [63–65]. It has been also suggested that glycolysis-driven changes in the intracellular pH may further favour *β*-catenin acetylation and nuclear translocation during NMP differentiation [14]. Glycolytic activity also feeds into the production of the building blocks necessary for glycosylation [66, 67]. Recently, it was proposed that 2-DG treatment affects mesoderm specification due to reduced glycosylation, based on the knowledge that several proteins involved in Wnt and Fgf signalling transduction are known to be glycosylated [20, 21, 68].

Besides post-translational modifications, glycolytic activity has also been shown to affect cellular localisation of glycolytic enzymes, thereby allowing them to perform non-canonical functions. For instance, translocation of Pfkl and Aldoa into the nucleus has been hypothesised to modulate Wnt signalling during somitogenesis [12]. Moreover, metabolites might also modulate protein activity by direct binding. Sentinel metabolites such as fructose-1,6-bisphosphate (FBP) whose concentration changes with glycolytic activity might be interesting targets for metabolite-protein allosteromes as suggested by Miyazawa and colleagues [12].

Most of the previous work has been focused on Wnt signalling and mesoderm specification. Currently, we know only little about how glycolytic activity modulates the Nodal signalling pathway and endoderm differentiation. Future work will be also important to identify actual changes in acetylation and glycosylation patterns of signalling components and functionally link them to their activity, as well as probe non-canonical glycolytic enzyme function and possible metabolite-protein interactions.

### Relevance for *in vivo* mouse gastrulation

The observation that exogenous Glc concentrations ranging from 0.02mM to 2.5mM have a dose-dependent effect on germ layer proportions, raises the question whether Glc levels within the embryonic environment fall within this regulatory range and whether glycolytic activity may indeed function as an instructive cue during *in vivo* gastrulation. Notably, measurements of Glc levels in uterine fluid suggest a concentration of approximately 0.6mM [27]. Interestingly, dose dependency is almost linear until 0.5mM and starts plateauing afterwards (Figure S3B). Hence, it is conceivable that differential expression of Glc transporters and glycolytic enzymes could account for spatial variations in glycolytic activity, thereby impacting cell fate decision making. This notion gains added significance when considered alongside recent findings that demonstrate the spatiotemporal coordination of Glc transporter expression during mouse gastrula development [20].

Our study provides new insights into the interplay between metabolism and signalling pathways that coordinate the differentiation of pluripotent cells into the three germ layers. The demonstrated instructive role of exogenous Glc concentration on cell fate decision-making represents an initial stride toward establishing nutritional control of cell type composition in complex *in vitro* model systems. Future work in this emerging research field will further improve our understanding of how metabolism is integrated into cellular behaviour and how metabolic conditions affect embryonic development.

## Acknowledgments

We thank the Mesoscopic Imaging Facility (MIF) at the European Molecular Biology Laboratory (EMBL) for support. We further thank the EMBL Genomics Core Facility for sequencing or data processing and the EMBL Genome Biology Computational Support for data management and submission. We thank Joshua Frenster for advice on gastruloid dissociation and flow cytometry. We thank all members of the Trivedi and Ebisuya labs for insightful discussions and feedback throughout the project. We thank Idse Heemskerk, Katharina Sonnen, Charisios Tsiairis for feedback on the manuscript.

## Author contributions

Conceptualization: K.S.S., M.E., V.T.; Methodology: K.S.S., M.E., V.T.; Software: N.G.; Validation: K.S.S., E.H., L.S.P., Kr.A.; Formal analysis: K.S.S., L.S.P., Ke.A., P.P, M.O.L.; Investigation: K.S.S., E.H., L.S.P.; Writing – original draft: K.S., V.T.; Visualization: K.S.S., V.T.; Supervision: K.S.S., M.E., V.T.; Project administration: V.T.; Funding acquisition: K.S.S., M.E., V.T.

## Supplementary information

Materials and methods, Supplementary figures and Supplementary movies.

