## Supplementary material for "Metabolic control of germ layer proportions through regulation of Nodal and Wnt signalling": Stapornwongkul_SuppInfo.pdf

**S4 Supplementary Movies** 10

**S5 Supplementary Figures** 10

[1]Tissue Biology and Disease Modelling European Molecular Biology Laboratory (EMBL) Barcelona, Spain. [2]Genomics Core Facility, European Molecular Biology Laboratory (EMBL) Heidelberg, Germany. [3]Developmental Biology, European Molecular Biology Laboratory (EMBL) Heidelberg, Germany. [4]Cluster of Excellence Physics of Life, TU Dresden, Dresden, Germany. [5]Max Planck Institute of Molecular Cell Biology and Genetics, Dresden, Germany. [6]These authors contributed equally.*

S4 **Wnt signalling activation can partially rescue gastruloid development but not glycolytic activity.** 14

### List of Tables

S1 Key Resource Table . . . . . . . . . . . . . . . . . . . . . . . . . . . . . . . . . . . . . . 9

### S1 Resource Availability

**Lead contact**

**Materials availability**
This study did not generate new unique reagents.

**Data and code availability**
• RNA-seq data have been deposited at BioStudies and are publicly available as of the date of publication. Accession numbers are listed in the key resources table. Microscopy data reported in this paper will be shared by the lead contact upon request.
• All original code has been deposited at GitHub and is publicly available as of the date of publication. DOIs are listed in the key resources table.
• Any additional information required to re-analyze the data reported in this paper is available from the lead contact upon request.
• The Python code used to perform image analysis is publicly available and deposited at: https://git.embl.de/grp-mif/image-analysis/gastr-volume-hcr.

#### S2 Materials and methods

##### S2.1 Mouse embryonic stem cell culture

E14 T/Bra::GFP mESCs (Fehling et al., 2003) (kindly shared by the Martinez Arias lab) were cultured as previously described (Anlas et al., 2021). T/Bra::GFP mESCs were maintained in ESL medium on 0.1% gelatin-coated (Millipore, #ES-006-B) tissue culture-treated 25cm$^2$ flasks (Corning, CLS430639) in a humidified incubator (37°C , 5% $CO_2$). ESL (KnockOut DMEM (Life technologies, 10829-018) supplemented with 10% ES tested fetal bovine serum (Life technologies, 16141-079), 1x Non-essential amino acids (Life technologies, 11140-035), 50 U/ml Pen/Strep (Life technologies, 15140-122), 1x GlutaMax (Life technologies, 35050-0610), 50$\mu$M 2-Mercaptoethanol (Life technologies, 31350-010) and 1000U/ml leukemia inhibitory factor (Life technologies, PMC9484)) was prepared and provided by the Tissue Engineering Unit of the Centre for Genomic Regulation. Cells were passaged every 2-3 days and the ESL was exchanged on alternating days. Cells were cultured up to 50-80% confluency prior to passaging or experimental use.

##### S2.2 Gastruloid generation and culture

Gastruloids were generated as previously described (Anlas et al., 2021) with following modifications: In none of the experiments, we have treated gastruloids with CHIR between 48-72hpa as indicated in the original protocol. While the CHIR treatment is beneficial for gastruloid elongation, robust symmetry breaking and elongation is achieved in the absence of CHIR addition when E14 T/Bra::GFP mESCs were maintained in ESL.

##### S2.3 Inhibitor and signalling agonist treatments

Gastruloids were made by aggregating 300 cells in 40$\mu$l N2B27 (Ndiff227, Takara, Y40002). At 24hpa selected inhibitors and signalling agonists were applied for 24h (till 48hpa) as 40$\mu$l 2x solutions in N2B27. Following final concentrations were used: glycolysis inhibition with 6mM 2-DG (stock solution: 2.5M 2-DG in dd$H_2O$, Sigma, D8375), OxPhos inhibition with 1mM $NaN_3$ (stock solution: 1M $NaN_3$ in dd$H_2O$, Sigma, S8032), Nodal signalling activation with 25ng/ml Act-A (stock solution: 100$\mu$g/ml Act-A in 2mM HCl, R&D Systems, #338-AC-010), Wnt signalling activation with 3$\mu$M CHIR (stock solution: 10mM CHIR in DMSO, Sigma, #SML1046), Fgf signalling activation with 100ng/ml Fgf8 (Tebubio, AF-100-25). At 48hpa, inhibitors and signalling agonists were washed out by adding 150$\mu$l of fresh N2B27, removing 180$\mu$l medium and adding another 150$\mu$l of fresh N2B27. At 72hpa, 140$\mu$l medium was removed and 150$\mu$l of fresh N2B27 medium was added.

##### S2.4 Washout and glucose concentration experiments

For Glucose (Glc) titration and washout experiments, homemade N2B27 was prepared by mixing equal parts of SILAC DMEM/F12 (Gibco, A2494301) and Neurobasal medium without glucose or pyruvate (Gibco, A2477501) and adding 0.5x N2 supplement (Gibco, 17502001), 0.5x B27 supplement (Gibco, 17504001), 1x GlutaMAX (Gibco, 35050061), 1mM sodium pyruvate (Gibco, 11360-070), 1x MEM non-essential amino acids (Gibco, 11140-050), 0.1mM 2-mercaptoethanol (Gibco, 31350-010) and respective concentrations of Glc (Agilent, 103577-100).
If not stated otherwise, 350 cells were aggregated in 40$\mu$l 0.5mM Glc N2B27. For the Glc washout experiments, gastruloids were repeatedly washed in Glc-free medium. At 24 hpa, 170$\mu$l Glc-free N2B27 was added, and subsequently removed. This was repeated twice resulting in a final estimated Glc concentration of below 0.005mM Glc. This was denoted as "no Glc" condition. At 48hpa, medium was changed back to 0.5mM Glc with a final volume of 150$\mu$l.
For Glc concentration experiments, gastruloids were aggregated and cultured in the media with respective Glc concentration. At 24hpa, 40$\mu$l of fresh medium with the respective Glc concentration was added. At 48hpa, 110$\mu$l of fresh medium with the respective Glc concentration was added.

#### S2.5 HCR staining and imaging

In situ mRNA detection was performed with the $HCR^{TM}$ RNA-FISH Technology (Molecular Instruments). Gastruloids were fixed for 40min in 4% paraformaldehyde (ChemCruz, SC-281692) and subsequently washed 3 times in PBS without $Mg^{2+}$/$Ca^{2+}$ (PBS-/-, Sigma, D8537). Gastruloids were then dehydrated in MeOH and subsequently stored in fresh MeOH at -20°C for overnight or longer. Gastruloids were rehydrated by sequential washes in 75% MeOH, 50% MeOH and 25% MeOH in PBS-/- supplemented with 0.1% Tween20 (Sigma, P9416) (PBST) and finally washed five more times in PBST. Gastruloids were prehybridised in probe hybridisation buffer (PHB, Molecular Instruments, $HCR^{TM}$ RNA-FISH Probe Set) for at least 30min at 37°C. Pre-PHB was then replaced with PHB containing 0.4pmol/100$\mu$l of each probe of interest (Molecular Instruments, commercially designed probes using accession# as indicated in Key resource table) and samples were incubated at 37°C overnight. Samples were then washed four times in probe wash buffer (PWB, Molecular Instruments) for 15min each at 37°C and two times in 5x sodium chloride sodium citrate with 0.1% Tween (5x SSCT, prepared in-house) for 5min at RT, each. Next, gastruloids were pre-incubated in amplification buffer (Molecular Instruments) for 30min at RT. In the meantime, all h1 and h2 hairpins (Molecular Instruments) of one reaction were separately mixed, heated to 95°C for 90s and cooled for 30min in the dark at RT. Then, h1 and h2 hairpins of each reaction mix were combined in amplification buffer at RT to a final concentration of 60nM per hairpin. The amplification buffer containing the hairpins was added to the samples after removing the pre-amplification buffer and samples were incubated overnight in the dark at RT. On the last day, five washes in 5x SSCT were conducted at RT: 2x 5min, 2x 30min and 1x 5min. Samples were stored at 4°C in the dark for up to 2 weeks before imaging on an Opera Phenix HSC system (PerkinElmer).

#### S2.6 Wide field imaging and time lapse microscopy

For live imaging, ULA 96-well plates containing gastruloids were transferred into the Opera Phenix HSC system (Perkin Elmer) and snapshots taken every 24h at 37°C and 5% $CO_2$ using a 10x air (NA 0.3) objective in wide field mode. The same settings were applied for time lapse imaging of gastruloids in ULA 96-well plates covered with MicroClime environmental microplate lids (Labcyte, LLS-0310) filled up with sterile $H_2O$ or PBS (Thermo Fisher Scientific) to avoid medium evaporation. Images were taken every 5min for several days with medium changes every 24h. For imaging of fixed gastruloid samples on the Opera Phenix HSC system, stained gastruloids were transferred into individual wells of flat-bottom CellCarrier 96-well (Perkin Elmer, 6007008) in SSCT. Images were acquired in confocal mode using a 20x water objective.

#### S2.7 RNA-seq sample preparation

We performed bulk RNA-seq on single gastruloids (2-DG treated between 24-48hpa and control, $n = 6$ each) at 48hpa (0h after 2-DG treatment) and at 96hpa (48h after 2-DG treatment). RNA was extracted from fresh single gastruloids using the RNeasy Micro Kit (QIAGEN). RNA quality control was performed with the Agilent RNA 6000 Pico Kit (Agilent, 5067-1513) on the Bioanalyzer system (Agilent).

#### S2.8 RNA-seq library preparation and sequencing

The extracted RNA was concentrated using a vacuum concentrator centrifuge (SpeedVac, Eppendorf, EP5305000100-1EA) to a final volume of 3$\mu$l. Then 1$\mu$l of 10mM dNTPs (Kappa, KK1017) and 1$\mu$l of oligo-dT primer (5'-AAGCAGTGGTATCAACGCAGAGTACT30VN-3, custom from IDT) were added to the samples. Then full-length cDNA sequencing libraries were prepared following a modified version of the SMART-Seq2 protocol using SuperScript IV RT (SSRT, ThermoFisher 18090050) and the tagmentation procedure previously described (Hennig et al., 2018; Picelli, 2019). Briefly, a reverse transcription reaction was performed with the following mixture per sample and cycling conditions: 2$\mu$l SSRT IV 5x buffer (ThermoFisher 18090050), 0.5$\mu$l 100mM DTT (ThermoFisher 18090050), 2$\mu$l 5M betaine (Merck, B0300-5VL), 0.1$\mu$l 1M MgCl2 (Ambion, AM9530G), 0.25$\mu$l 40 U/$\mu$l RNAse inhibitor (Clontech Takara, 2313A), 0.25$\mu$l SSRT IV (ThermoFisher 18090050), 0.1$\mu$l 100$\mu$M TSO primer (5'-AAGCAGTGGTATCAACGCAGAGTACATrGrG+G-3') and 1.15$\mu$l RNase-free $H_2O$ (Ambion,

AM9916) at 52°C for 15min, followed by an incubation at 80°C 10min. The resulting cDNA was amplified adding 12.5$\mu$l of KAPA HiFI HotStart Ready Mix 2x (Roche, 07958935001), ISPCR primer at 5$\mu$M (5'-AAGCAGTGGTATCAACGCAGAGT-3') and 2.30$\mu$l of $H_2O$. The cycling conditions were 98°C for 3min, followed by 18 cycles of 98°C for 20s, 67°C for 15s and 72°C for 6min and a final incubation at 72°C for 5min. After amplification, individual libraries were size selected using magnetic SPRI bead purification (Beckman coultier, B23319) (0.6x) omitting the ethanol wash steps and using an elution volume of 13$\mu$l $H_2O$. For the next steps, the clean cDNA was normalized to 0.2ng/$\mu$l. In short, 1.25$\mu$l of each sample was taken into a tagmentation reaction containing 1.25$\mu$l of Dimethylformamide (Merck, 227056), 1.25$\mu$l of tagmentation buffer (40mM Tris-HCl pH 7.5, 40mM MgCl2) and 1.25$\mu$l of an in-house generated and purified Tn5 (previously prepared as described by Hennig et al. 2018 and diluted to a 1:300 ratio with water). The mixture was incubated at 55°C for 3min. After that, 1.25$\mu$l of 0.2% SDS (Merck, 71736) was added to stop the tagmentation reaction followed by 5min incubation at room temperature. The resulting fragments were then amplified by PCR using 6.75$\mu$l of KAPA 2x HiFi master mix, 0.75$\mu$l of Dimethyl sulfoxide (Merck, D8418-50ML) and 2.5$\mu$l of dual indexed primers. Cycling conditions were 72°C for 3min and 95°C for 30s, followed by 12 cycles of 98°C for 20s, 58°C for 15s, 72°C for 20s and a final incubation at 72°C for 3min. Final libraries were size selected with the magnetic SPRI beads (0.8x) and were quantified using Qubit HS assay (Thermo Fisher Scientific, Q33231). A Bioanalyzer system (Agilent, 5067-4626) was used to assess library sizes and fragment distribution. Finally, libraries were pooled equimolarly and the final pool underwent an extra round of size selection using magnetic SPRI beads at a 0.7x ratio. Samples were single end sequenced as a 24-plex on Illumina NextSeq2000 using a P2 100 cycles kit (Illumina, 20046811).

#### S2.9 Seahorse metabolic rate assay

To mimic a differentiation timeline similar to gastruloid development, mESCs were seeded at a density of $1.67x10^5$ cells/T25 flask in ESL medium. The following day cells were washed twice in PBS (Sigma, D8662) before adding N2B27. After 24h, medium was replaced with N2B27 supplemented with metabolic inhibitors and/or signalling agonists, as indicated for the different experiments. After 24h of treatment, cells were dissociated with trypsin-EDTA (Thermo Fisher Scientific) for 1 min at 37°C , harvested in their respective culture media, washed and resuspended in N2B27 medium +/- inhibitors/signalling agonists. 80,000cells/well in 100$\mu$l medium were seeded into fibronectin-coated Seahorse cell culture plates (Agilent, 100777-004) and left at RT for 15 min to ensure equal distribution of cells within all microwells. Plates were then transferred into a 37°C incubator for 2h for complete attachment. Afterwards, cells were washed carefully in Seahorse XF DMEM medium (Agilent, 103575-100) supplemented with 1mM pyruvate (Agilent, 103578-100), 2mM glutamine (Agilent, 103579-100) and the respective concentration of Glc (Agilent, 103577-100) +/- inhibitors/signalling agonists. If not indicated otherwise, 10mM Glc were used as a standard concentration. During the washing steps, 950$\mu$l Seahorse medium were added to the attached cells, 1000$\mu$l removed, another 450$\mu$l added and removed and a final 450$\mu$l added onto the cells (final volume 500$\mu$l). 500$\mu$l Seahorse medium was added to each of the 4 background wells. Cells were then incubated at 37°C without $CO_2$ for 45 min. The Seahorse cartridge was hydrated overnight and loaded with components of the real-time ATP rate assay (Agilent, 103592-100) as indicated in the manual (final inhibitor concentration: 1.5$\mu$M oligomycin, 0.5$\mu$M rotenone and 0.5$\mu$M antimycin A). All samples were run with at least 4 technical replicates in a Seahorse XFe24 Analyzer (Agilent).

#### S2.10 Flow cytometry

To determine how size and Glc concentration affect the percentage of T/Bra positive cells, gastruloids from 4 different initial cell numbers (350, 700, 1050 and 1400 cells) were grown at 0.02mM, 0.1mM, 0.5mM and 2.5mM Glc until 48hpa. 24 gastruloids of each condition (48 for initial cell number of 350 cells) were pooled and washed in PBS-/- before adding 50$\mu$l of accutase (Gibco, A1110501). After 5 min of incubation in accutase, gastruloids were dissociated by adding flow buffer (20% N2B27 with 0.02mM Glc and 2.5mM EDTA in PBS-/-) and gentle up and down pipetting. Cells were then pelleted at 170xg for 3min and resuspended in 200$\mu$l flow buffer. Flow cytometry was performed on BD LSRFortessa Cell Analyzer (BD Biosciences).

### S3 Quantification and Statistical Analysis

#### S3.1 Gastruloid size and Bra/T intensities

Mean T/Bra::GFP intensity (with background subtraction) and gastruloid size at 48hpa were quantified using Morgana v.0.1 (Gritti et al., 2021). In brief, after generation of segmentation masks using the brightfield channel of widefield images obtained with the Opera Phenix HSC system (Perkin Elmer), Morgana was used to calculate the area and the mean T/Bra::GFP intensity (with background subtraction).

#### S3.2 Kymographs

Kymographs were generated using Morgana v.0.1 (Gritti et al., 2021). In brief, after generation of segmentation masks and calculation of AP fluorescence profiles for each gastruloid across the entire time-lapse, all experimental conditions to be compared were loaded simultaneously as separate groups of respective gastruloids. T/Bra::GFP fluorescence intensities along the thereby defined AP axis were normalized to the global maximum (yielding values between 0-1). The colour code denoting Bra fluorescence intensity was manually set between 0-0.83 (Figure 1B) and 0.05-0.95 (Figure 2D) normalized units to enable clear visual comparison of fluorescence intensities between conditions. Prior to normalization, background fluorescence as a result of the culture media or gastruloid tissue properties was removed either by subtracting the average fluorescence intensity of the non-mask area or by setting the minimum pixel intensity level found in the gastruloid mask to 0.

#### S3.3 Analysis of area of gene expression

The germ layer proportions were measured as the relative area of gene expression from the images of HCR stained gastruloids and obtained using custom written software using Python. Individual field of views (tiles) obtained from imaging on the Opera Phenix microscrope were flat-field corrected using the parameters obtained from the Harmony software (version 4.9), and full-well views were generated stitching the tiles with the Grid/Collection Stitching plugin of FIJI (Preibisch et al., 2009). Semantic segmentation was performed with a pixel classification network in the ilastik software v. 1.4 (Berg et al., 2019) to classify pixels belonging to gastruloids or background using the bright field images. To perform instance segmentation, that is, identification of individual gastruloids in each well, we eroded the ilastik masks to separate individual gastruloids, removed objects smaller than $1000 \mu m^2$ and computed a distance transform of the masks using the Python package scipy (Virtanen et al., 2020). The locations of the max values of the distance transform image were used as seeds of a watershed segmentation using scikit-image (van der Walt et al., 2014). Finally, using the histogram of pixel values from all gastruloid masks in the image, we obtained a threshold value for each fluorescent channel using the otsu algorithm, and computed gene expression masks. The relative gene expression area was computed as the total number of bright pixels in each gastruloid divided by the total area of the same gastruloid. To avoid bias in threshold computation under conditions in which no gene expression was found, in each experiment, we always had one control condition in which the otsu thresholds were computed, and subsequently the same thresholds were applied to all other metabolic conditions in the same experiment. When the otsu algorithm failed to generate the correct thresholds, we set such thresholds manually by visual inspection. The Python code used to perform image analysis is publicly available and deposited at: https://git.embl.de/grp-mif/image-analysis/gastr-volume-hcr.

#### S3.4 RNA-seq analysis

We have aligned the sequencing reads to GRCm39 reference (version 107) using STAR Aligner, version: 2.7.9a; Link: https://github.com/alexdobin/STAR (Dobin et al., 2013). Aligned reads were counted using HTSeq-count, version 2.0.2; Link: https://htseq.readthedocs.io/en/master/ (Anders et al., 2015; Putri et al., 2022). Differential gene expression was assessed with DESeq2 (Love et al., 2014). $\text{Log}_2$ fold changes between vehicle and 2-DG treated gastruloids were shrunk using rlog function. RNA-seq data have been deposited in the BioStudies, and is available at the E-MTAB-13543 accession number.

#### S3.5 Seahorse metabolic rate analysis

The Wave online app (Agilent) was used for analysis & Graphpad Prism (Graphpad) for plot generation.

#### S3.6 Flow cytometry analysis

Data was analysed using the FlowJo software (BD Life Sciences).

#### S3.7 Key Resource Table

See next page

| Reagent or Resource | Source | Identifier |
|---|---|---|
| **Chemicals, peptides, and recombinant proteins** | | |
| Ndiff | Takara | Cat# 40002 |
| 2-Deoxyglucose | Sigma | Cat# D8375 |
| $NaN_3$ | Sigma | Cat# S8032 |
| Activin-A | Bio-Techne | Cat# 338-AC |
| CHIR99021 | Merck | Cat# SML1046 |
| Fgf8 | T ebubio | Cat# AF-100-25 |
| Glucose | Agilent | Cat#103577-100 |
| **Critical commercial assays** | | |
| HCR$^{TM}$ RNA-FISH Bundle | Molecular Instruments | https://store.molecularinstruments.com/ |
| Seahorse XF Real-Time ATP Rate Assay Kit | Agilent | Cat#103592-100 |
| **Deposited data** | | |
| Single gastruloid RNA sequencing data | This paper | BioStudies E-MTAB-13543 |
| **Oligonucleotides** | | |
| HCR probe: Aldh1a2 | Molecular Instruments | Acc# NM_009022 |
| HCR probe: Eomes | Molecular Instruments | Acc# NM_010136.3 |
| HCR probe: Gata6 | Molecular Instruments | Acc# NM_010258 |
| HCR probe: Meox1 | Molecular Instruments | Acc# NM_010791 |
| HCR probe: Nanog | Molecular Instruments | Acc# NM_001289828.1 |
| HCR probe: Sox1 | Molecular Instruments | Acc# NM_009233.3 |
| HCR probe: Sox17 | Molecular Instruments | Acc# NM_011441.5 |
| HCR probe: Sox2 | Molecular Instruments | Acc# NM_011443.4 |
| HCR probe: T/Bra | Molecular Instruments | Acc# NM_009309.2 |
| **Software and algorithms** | | |
| MOrgAna v.0.1 | https://github.com/LabTrivedi/MOrgAna/ | Gritti et al., 2021 |
| FIJI/Image J | NIH | https://imagej.net/Fiji |
| Python | Python Software Foundation | https://www.python.org |
| ilastik v.1.4 | https://www.ilastik.org/ | Berg et al., 2019 |
| Python script for analysis of gene expression areas | This paper | https://git.embl.de/grp-mif/image-analysis/gastr-volume-hcr |
| FlowJo™ | BD Life Sciences | https://www.flowjo.com |
| Wave online app | Agilent | https://www.seahorseanalytics.agilent.com/ |
| Graphpad Prism | Graphpad Software | https://www.graphpad.com/ |
| **Other** | | |
| 96-well ULA plates | Greiner | Cat#650970 |
| Flat-bottom CellCarrier 96 ultra plate | PerkinElmer | Cat#6007008 |
| MicroClime environmental microplate lid | Labcyte | Cat#LLS-0310 |
| Seahorse cell culture plates | Agilent | Cat# 100777-004 |

**Table S1.** Key Resource Table.

#### S4 Supplementary Movies

**Movie S1: Symmetry breaking and elongation in a control gastruloid**
Timelapse imaging of control T/Bra::GFP gastruloid (24-96hpa). GFP shown in a fire lookup table.

**Movie S2: 2-DG treated gastruloid does not upregulate T/Bra::GFP expression**
Timelapse imaging of 2-DG treated T/Bra::GFP gastruloid (24-96hpa, 2-DG treatment: 24-48hpa). GFP shown in a fire lookup table.

#### S5 Supplementary Figures

See next page

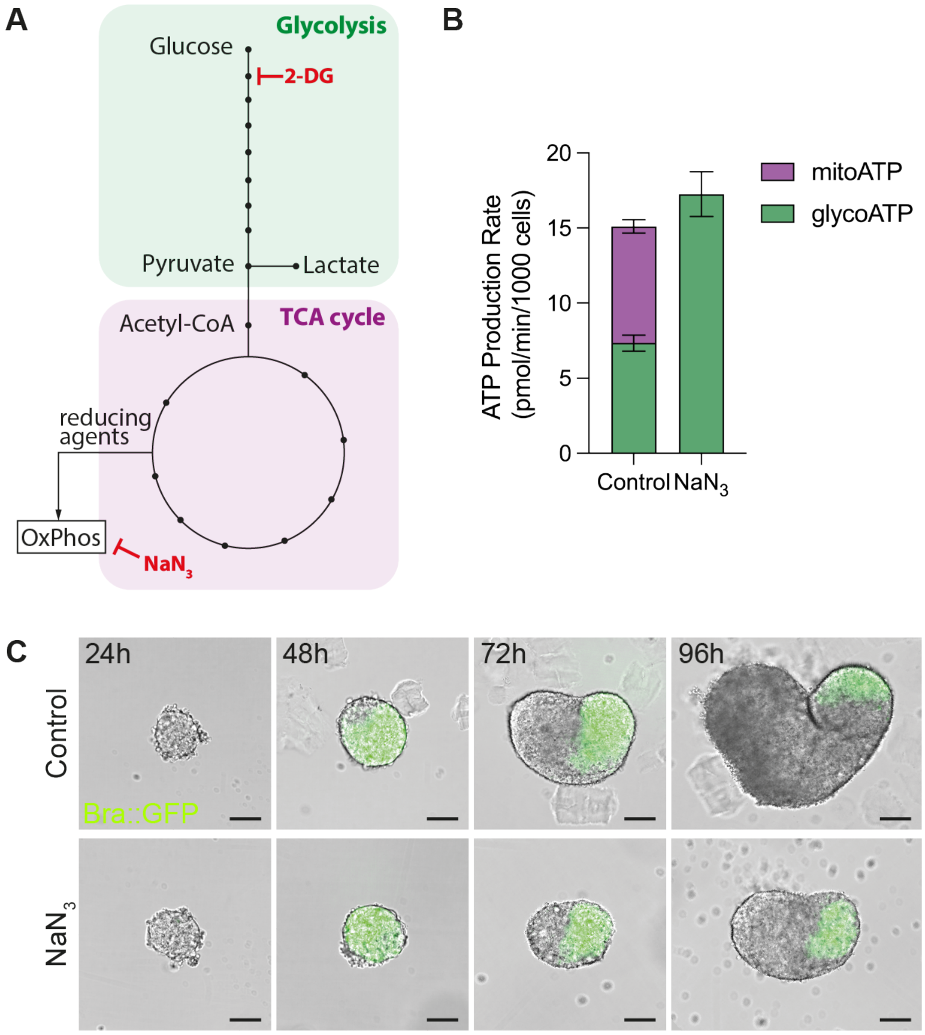

**Figure S1. Inhibition of oxidative phosphorylation affects growth but not T/Bra expression.**

**(A)** Simplified schematic of the central carbon metabolism. Aerobic glycolysis (green) results in the production of lactate from glucose. This process yields comparatively little ATP but allows the rapid production of building blocks needed for proliferation and growth. Alternatively, pyruvate can be shuttled into mitochondria to produce acetyl-CoA. Acetyl-CoA enters the tricarboxylic acid (TCA) cycle (purple) which generates reducing agents needed for the production of large amounts of ATP during OxPhos. Please note that the TCA cycle can be also fueled by amino and fatty acids. 2-DG acts as a competitive inhibitor and blocks glycolytic activity on the level of hexokinase and glucose-6-phosphate isomerase. Therefore, lactate production is strongly reduced during 2-DG treatment. Sodium azide ($NaN_3$) blocks OxPhos and TCA cycle activity by inhibiting Complex IV of the electron transport chain.

**(B)** Contribution of glycolysis and OxPhos to total ATP production rate (glycoATP and mitoATP) in control cells and $NaN_3$ treated cells. Cells were cultured in N2B27 for 48h prior Seahorse measurement. Treatment was performed between 24-48h. $N_{exp}$ =3, at least 4 replicates in each experiment.

**(C)** Gastruloids treated with 1mM $NaN_3$ between 24-48hpa show growth inhibition similar to the 2-DG treated gastruloids but no loss of T/Bra::GFP upregulation and symmetry breaking. $N_{exp}$ =3, control: $n$ =83/84 (control), $n$ =75/84 ($NaN_3$). Scale bars: 100$\mu$m.

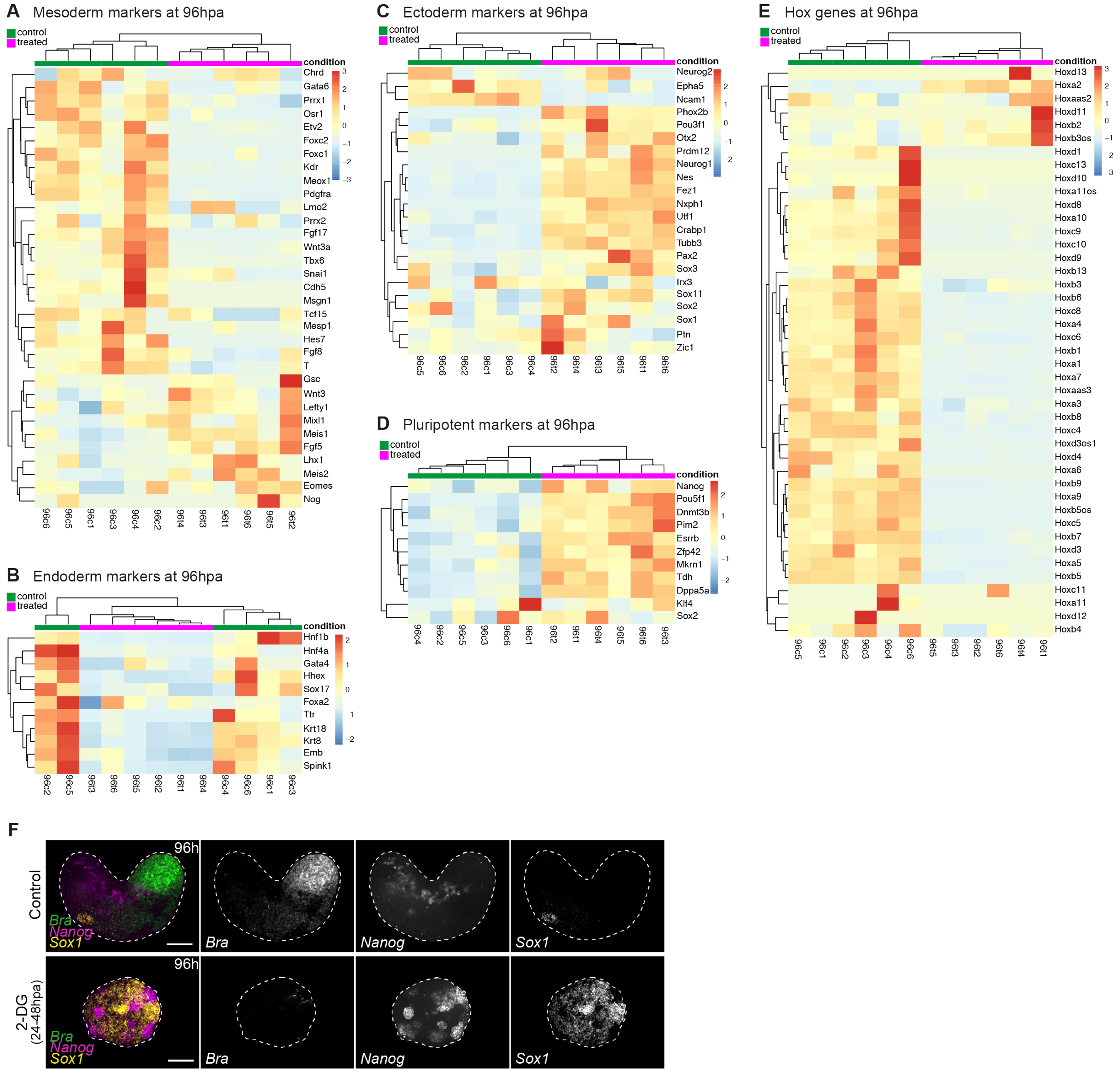

**Figure S2. Glycolysis inhibition results in an increased proportion of neuroectodermal and pluripotent cells.**
**(A-E)** Heatmaps describing relative transcript levels of selected fate cell-fate markers (A-D) and Hox genes (E) based on RNA-seq. Heatmap rows are normalised per gene. Lower and higher expression in blue and red, respectively. Each column represents a single gastruloid with applied hierarchical clustering.
**(F)** HCR stainings of control and 2-DG treated gastruloids. *T/Bra*: PS/early mesoderm, *Nanog*: pluripotent cells, *Sox1*: neural progenitors. $n$ =10 (control), $n$ =17 (2-DG). Maximum $z-$projection of 6 confocal slices is shown. Scale bar equals 100$\mu$m.

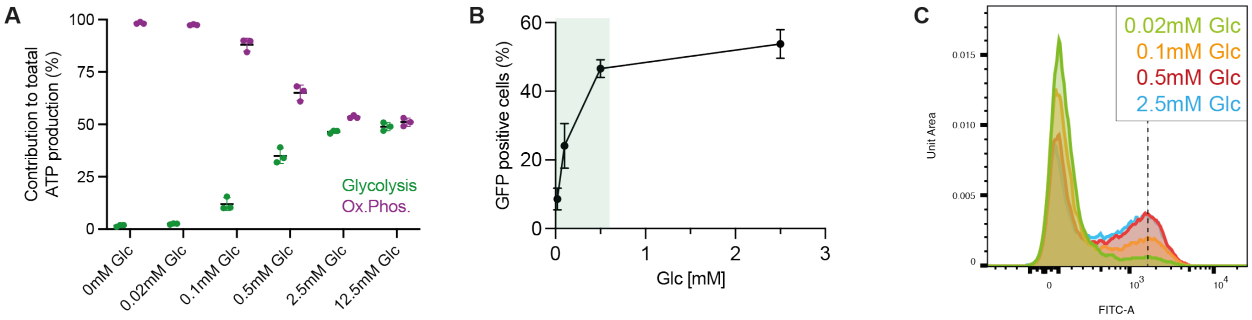

**Figure S3. Modulation of glycolytic activity and number of T/Bra::GFP expressing cells with exogenous glucose levels.** **(A)** Contribution of glycolysis and OxPhos to total ATP production dependent on exogenous Glc concentration. $N_{exp} = 3$, 4 replicates in each experiment.

**(B)** Mean percentage of GFP$^+$ cells in T/Bra::GFP gastruloids at 48hpa which were generated by aggregating 350, 700, 1050 and 1400 cells and cultured in different Glc concentrations (same data as in Figure 2C but on a linear scale). Green shaded area: $0 - 0.6$ mM Glc.

**(C)** Histogram of cells from T/Bra::GFP gastruloids grown at different Glc concentrations. While the proportion of GFP$^+$ cells changes with Glc concentration, typical GFP intensity in GFP$^+$ cells is not shifted (indicated by dashed horizontal line).

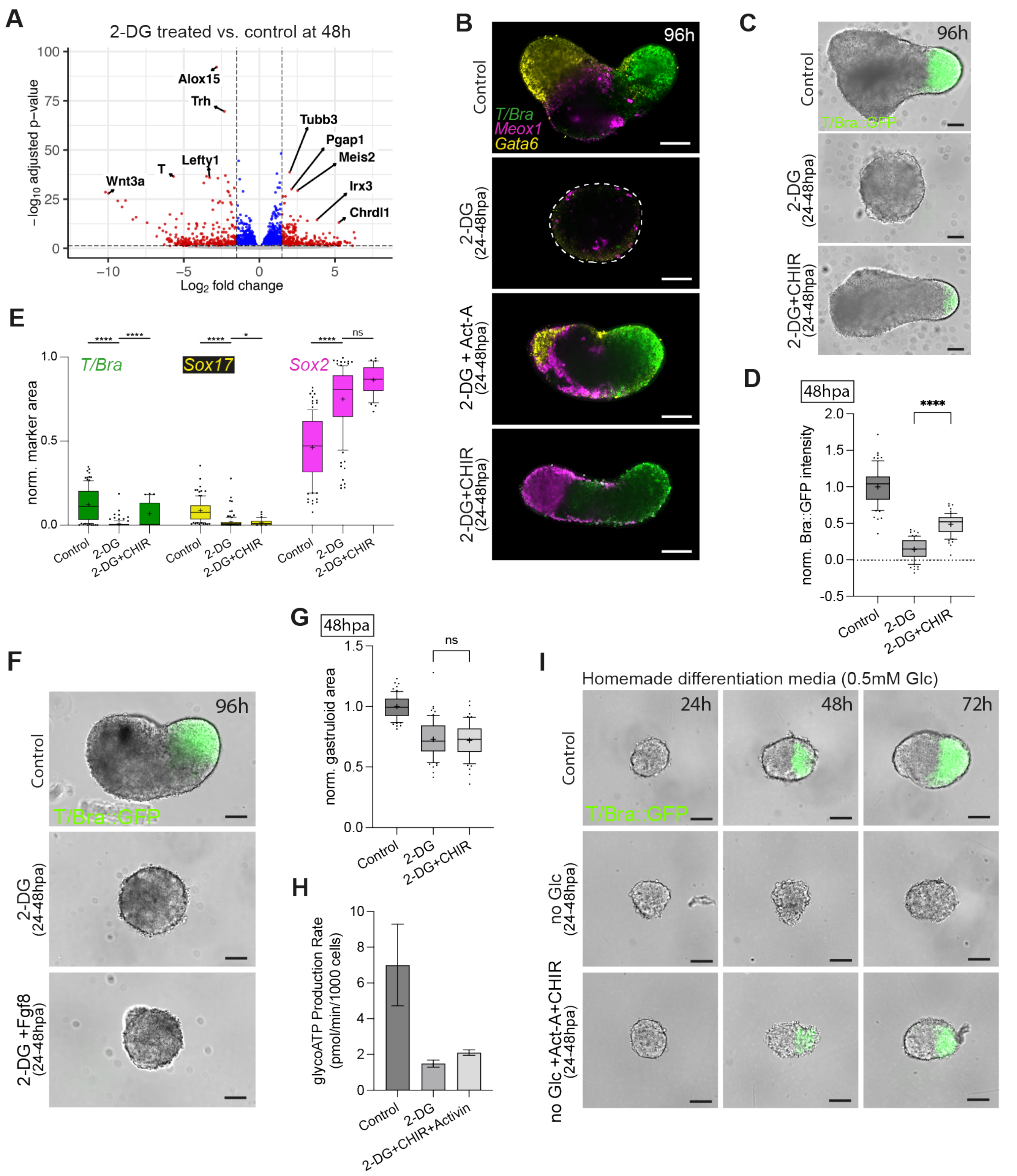

**Figure S4. Wnt signalling activation can partially rescue gastruloid development but not glycolytic activity.**

**(A)** Volcano plot describing differentially expressed genes between 2-DG treated and control gastruloids at 48hpa based on RNA-seq. Red dots describe significantly differentially expressed genes that pass thresholds for log2FC = +/-1.5 (vertical lines) and adjusted p value = 0.05 (horizontal line). $n$ =6 for both conditions.

**(B)** HCR stainings of control, 2-DG treatment, Nodal (2-DG+Act-A) and Wnt (2-DG+CHIR) signalling rescue. *T/Bra* (green), *Meox1*: paraxial mesoderm (magenta), *Gata6*: cardiac mesoderm/endoderm (yellow). A single confocal section is shown. $n = 21$ (control), $n = 17$ (2-DG), $n = 14$ (2-DG+Act-A), $n = 18$ (2-DG+CHIR). Scale bar 100$\mu$m.

**(C)** Gastruloids at 96hpa under control conditions, glycolysis inhibition or Wnt signalling rescue condition. $N_{exp} = 4$, symmetry breaking: $n = 82/85$ (control), $n = 0/88$ (2-DG), $n = 50/82$ (2-DG+CHIR); elongation: $n = 81/85$ (control), $n = 0/88$ (2-DG), $n = 28/82$ (2-DG+CHIR). Scale bar 100$\mu$m.

**(D)** T/Bra::GFP intensity of gastruloids at 48hpa for different treatment conditions. Background was subtracted and intensities were normalised to mean control intensity. Line in box plot indicates median, + indicates mean, points beyond whiskers (10-90 percentiles) are drawn as individual points. $N_{exp} = 3$, $n = 62$ (control), $n = 63$ (2-DG), $n = 59$ (2-DG+CHIR). ****$p < 0.0001$ for t-test.

**(E)** Area quantifications of germ layer markers using average $z$-projections of HCR stainings. Area was normalised to total gastruloid area. Line in box plot indicates median, + indicates mean, points beyond whiskers (10-90 percentiles) are drawn as individual points. $N_{exp} = 3$, $n = 107$ (control), $n = 104$ (2-DG), $n = 33$ (2-DG+CHIR). ****$p < 0.0001$, *$0.01 < p < 0.05$ for t-test.

**(F)** Gastruloids at 96hpa under control conditions, glycolysis inhibition or Fgf signalling rescue condition. $N_{exp} = 3$, symmetry breaking and elongation: $n = 60/64$ (control), $n = 0/64$ (2-DG), $n = 0/63$ (2-DG+Fgf8).

**(G)** Gastruloid area at 48hpa for different treatment conditions. Area was normalised to mean control area. Line in box plot indicates median, + indicates mean, points beyond whiskers (10-90 percentiles) are drawn as individual points. $N_{exp} = 3$, $n = 62$ (control), $n = 63$ (2-DG), $n = 59$ (2-DG+CHIR). Scale bars 100$\mu$m.

**(H)** Mean glycoATP production rate of cells cultured for 48h in differentiation medium. Inhibitor and growth factor treatment was started 24h prior to measurements. $N_{exp} = 3$, error bars indicate SD.

**(I)** Time course of gastruloids developing in homemade differentiation media containing 0.5mM Glc. Control gastruloids express T/Bra::GFP and break symmetry. Removing Glc between 24-48hpa results in loss of T/Bra::GFP expression. Addition of Act-A and CHIR in no Glc condition (24-48hpa) can rescue T/Bra::GFP expression and symmetry breaking. $N_{exp} = 2$, symmetry breaking: $n = 34/39$ (control), $n = 0/40$ (no Glc), $n = 40/40$ (no Glc+CHIR+Act-A). Scale bars 100$\mu$m.
